# Peripheral Serum Metabolomic Profiles Inform Antecedent Central Cognitive Impairment in Older Adults

**DOI:** 10.1101/837989

**Authors:** Jingye Wang, Runmin Wei, Guoxiang Xie, Matthias Arnold, Gregory Louie, Siamak MahmoudianDehkordi, Colette Blach, Rebecca Baillie, Xianlin Han, Philip L. De Jager, David A. Bennett, Rima Kaddurah-Daouk, Wei Jia, for the Alzheimer’s Disease Metabolomics Consortium

**Affiliations:** University of Hawaii Cancer Center, Honolulu, HI, USA; Department of Molecular Biosciences and Bioengineering, University of Hawaii at Manoa, Honolulu, HI, USA; Department of Psychiatry and Behavioral Sciences, Duke University, Durham, NC, USA; Institute of Bioinformatics and Systems Biology, Helmholtz Zentrum München, German Research Center for Environmental Health, Neuherberg, Germany; Duke Molecular Physiology Institute, Duke University, Durham, NC, USA; Rosa & Co LLC, San Carlos, CA, USA; University of Texas Health Science Center at San Antonio, San Antonio, TX, USA; Columbia University College of Physicians and Surgeons Department of Neurology, Center for Translational & Computational Neuroimmunology, New York, NY, USA; Rush Alzheimer’s Disease Center, Rush University Medical Center, Chicago, IL, USA; Institute of Brain Sciences, Duke University, Durham, NC, USA; Department of Medicine, Duke University, Durham, NC, USA

**Author notes:** Correspondence to: Wei Jia, University of Hawaii Cancer Center, 701 Ilalo Street, Honolulu, HI 96813, USA., Rima Kaddurah-Daouk, Duke University, Durham, NC, USA.

## Abstract

The incidence of Alzheimer’s disease (AD) increases with age and is a significant cause of worldwide morbidity and mortality. However, the metabolic perturbations behind the onset of AD remains unclear. In this study, we performed metabolite profiling in both brain (n = 109) and matching serum samples (n = 566) to identify differentially expressed metabolites and metabolic pathways associated with neuropathology and cognitive performance and to identify individuals at high risk of developing cognitive impairment. The abundances of six metabolites, GLCA, petroselinic acid, linoleic acid, myristic acid, palmitic acid, and palmitoleic acid as well as the DCA/CA ratio, along with the dysregulation scores of three metabolic pathways, primary bile acid biosynthesis, fatty acid biosynthesis, and biosynthesis of unsaturated fatty acids showed significant differences in diagnostic groups across both brain and serum (*P*-value < 0.05).Significant associations were observed between the levels of differential metabolites/pathways and cognitive performance, neurofibrillary tangles, and neuritic plaque burden. Metabolites abundances and personalized metabolic pathways scores were used to derive machine learning models that could be used to differentiate cognitively impaired persons from those without cognitive impairment (median of AUC = 0.772 for the metabolite level model; median of AUC = 0.731 for the pathway level model). Utilizing these two models on the entire baseline control group, we identified those who experienced cognitive decline (AUC = 0.804, sensitivity = 0.722, specificity = 0.749 for the metabolite level model; AUC = 0.778, sensitivity = 0.633, specificity = 0.825 for the pathway level model) and demonstrated their pre-AD onset prediction potentials. Our study provides a proof-of-concept that it is possible to discriminate antecedent cognitive impairment in older adults before the onset of overt clinical symptoms using metabolomics. Our findings, if validated in future studies, could enable the earlier detection and intervention of cognitive impairment that may halt its progression.

## Introduction

Alzheimer’s disease (AD), one of the top 10 leading cause of death in the United States, is a challenge for health care systems resulting in increased economic burden as the population ages^1,2^. Currently, there is no therapy to prevent or slow AD progression, which may be due to the inability to detect AD before its progression into evident cognitive decline. Identification of early biomarkers associated with preclinical symptoms would allow the development of early intervention or preventive strategies^3^. Multiple neurochemical perturbations have been identified in AD, including amyloid precursor protein metabolism, phosphorylation of tau protein, and a wide range of metabolic perturbations^4^. Unfortunately, current biomarkers for early disease, including cerebrospinal fluid beta-amyloid and tau levels^5^, structural and functional magnetic resonance imaging^6^, the recent use of brain amyloid imaging^7^ or inflammaging^8^, and spinal fluid markers to track brain atrophy and deposition of cortical beta-amyloid and neurofibrillary tangles, are limited because they are either invasive, time-consuming, or expensive.

Recent studies have focused on obtaining biomarkers to identify features that differentiate subjects with cognitive impairment from elderly persons without cognitive impairment. Molecular markers sensitive to the underlying pathogenic factors will be highly relevant to early disease detection and facilitation of disease monitoring and treatment responses. Metabolomics is an unbiased approach to study small-molecule metabolites that offers hope for the discovery of more biomarkers for AD. This profiling technology has already been used to identify differential metabolites that can distinguish mild cognitive impairment (MCI) subjects who will develop AD from stable MCI^9^. Mounting evidence suggests that AD is closely accompanied with the abnormal bile acid (BA) metabolism^10–13^, free fatty acid (FFA) metabolism^14,15,26^, lipid metabolism^16,17^ and neurotransmitter metabolism^18^. BAs are increasingly recognized as important metabolic signaling molecules that modulate lipid, glucose, and energy metabolism^19^. More importantly, BAs in the brain act as neuroactive steroids^20^. Different classes of BAs can either inhibit or potentiate GABAα, or inhibit NMDA receptors while also exerting neuroprotective effects ^21,22^. Recent cross-sectional studies have shown differences in blood BAs in AD patients compared with non-cognitively impaired individuals^23,24^. Additionally, compared to control mice, researchers found an accumulation of FFAs in the hippocampus and cortex of AD mice^25^. Alterations of FFAs have been detected in postmortem AD brain tissues^14^ and serum samples^26^, which may indicate a switch to an alternative fuel source before the onset of clinical symptoms^27^. These observations give rise to the possibility that metabolic perturbations could presage the onset of cognitive impairment and therefore aid in the identification of individuals with higher risks by providing additional information to use with standard clinical markers. In this study, we performed a metabolite profiling in participants from a large, longitudinal cohort, with the goal of identifying metabolic changes as well as key metabolic pathways that might also serve as new predictors of future cognitive impairment in older adults.

## Materials and Methods

### Participants

Study data from the Rush Alzheimer’s Disease Center (RADC) Religious Orders Study (ROS) and Memory and Aging Project (ROS/MAP) were downloaded from the RADC Data Sharing Hub (https://www.radc.rush.edu/home.htm). A variety of clinical, demographic, and other variables are available through RADC (https://www.radc.rush.edu/docs/var/variables.htm), while omics datasets generated from biospecimens donated by ROS and MAP participants are made available through the AMP-AD Knowledge Portal. The ROS, which began in 1994, is a longitudinal clinical-pathologic cohort study of risk factors of cognitive decline and incident dementia run from the RADC that is comprised of individuals from religious communities (e.g., Catholic brothers, nuns, and priests) across the USA^28,29^. The Rush MAP, which began in 1997, includes participants from northeastern Illinois, USA with a broader range of socioeconomic status and life experiences^29^. Participants in both studies enroll without known dementia, agree to annual clinical evaluation, and organ donation. Both studies were approved by an Intuitional Review Board of Rush University Medical Center. All subjects signed an informed consent, an Anatomic Gift Act, and a repository consent to allow their biospecimens and data to be used for ancillary studies. Both studies are conducted by the same team of examiners and share a large common core of data collection at the item level to allow for efficient merging of data.

### Cognitive performance tests

Cognitive function was measured using a battery of 21 cognitive tests, 19 of which are used to measure five cognitive domains (i.e., episodic memory, working memory, semantic memory, perceptual orientation/visuospatial ability, and perceptual speed) (Table S1). Raw scores from each cognitive test are converted to z-scores and averaged to create a global cognitive performance measure using the mean and standard deviation at baseline (M=0, SD=1). For each domain, scores are created by averaging the z-scores which are based on mean and standard deviation from all baseline data. Additionally, the Mini-Mental State Examination (MMSE) served as a global measure of cognitive function.

### Clinical diagnoses

Medical conditions were documented via self-report and clinical evaluation. Clinical diagnoses each year were determined blinded to previously collected data. A three-step process for all clinical diagnoses starts with an actuarial decision tree based on the history of cognitive decline and impairment ratings in five cognitive domains based on cutoffs for 11 cognitive tests^30^. Then clinical judgment by a neuropsychologist for cognitive impairment and determination of dementia and its causes by a clinician (i.e., neurologist, geriatrician, second neuropsychologist, geriatric nurse practitioner)^31^. The diagnosis of AD follows the criteria of the National Institute of Neurological and Communicative Disorders and Stroke and the Alzheimer’s Disease and Related Disorders Association (NINCDS/ADRDA)^32^. Participants were categorized as a) AD, b) MCI if diagnosed with cognitive impairment by the neuropsychologist but not diagnosed dementia by the clinician^30^, and c) no cognitive impairment (NCI) if diagnosed without AD or MCI^33^. At the time of death, a final diagnosis is rendered by a clinician blinded to postmortem data using all clinical data.

### Neuropathology

Upon death, a postmortem neuropathological evaluation is implemented, and procedures follow those outlined by the pathologic dataset recommended by the National Alzheimer’s Disease Coordinating Center. Brains of deceased subjects were removed, weighed, cut into one cm-thick coronal slabs and stored. Each brain was examined for the neuropathological indices of common pathologies that contribute to cognitive impairment. The location, age, and volume of all macroscopic infarcts were recorded, and tissue was obtained for histological confirmation, in addition to the identification of microscopic infarctions, as previously described^34,35^. AD pathology was identified using the modified Bielschowsky silver stain technique and by use of the Consortium to Establish a Registry for Alzheimer’s Disease (CERAD) criteria^36^ and NIA-Reagan criteria^37^, while the assessment of neurofibrillary tangles was based on Braak criteria^38^ as described previously^39^. The CERAD score, a semi-quantitative measure of neuritic plaque burden, is made of 4 levels: 4 = no AD, 3 = possible AD, 2 = probable AD and 1 = definite AD. As recommended, CERAD scores were reclassified to a binary level: score 1∼2, score 3∼4. Seven categories of Braak stages were based on the region and severity of neurofibrillary tangles pathology.

### Metabolites quantification

Using targeted metabolomics protocols^40^ and profiling protocols^41^ established in previous studies, BAs were quantified by ultra-performance liquid chromatography triple quadrupole mass spectrometry (UPLC-TQMS) (Waters XEVO TQ-S, Milford, USA) and other metabolites were quantified by gas chromatography time-of-flight mass spectrometry (GC-TOFMS) (Leco Corporation, St Joseph, USA). Details are described in the Supporting Information.

### Statistical analysis

Stratifying by clinical diagnosis, continuous demographic variables were expressed as mean (standard deviation (SD)) and tested by Wilcoxon rank-sum test, while categorical demographic variables were expressed as n (percentage) and tested by Chi-square test. Missing values in quantitative metabolites due to limits of quantification were regarded as left-censored missing and imputed by GSimp^42,43^. Individual BA concentrations were normalized to the total BAs concentration (i.e., the proportion of total BAs). Metabolites were reported as median (25% quantile, 75% quantile) and tested by Wilcoxon rank-sum test. Due to the limited sample size of the AD group (11 participants) in serum samples, we combined MCI and AD participants into an aggregate group (MCI/AD) for the following data analysis. Log-transformed abundances were used in the following data analysis. We additionally generated 12 BA ratios based on the BA metabolic pathway topology.

To identify metabolites differentially expressed in participants with dementia and cognition, we used ordinal logistic regression to compare metabolites across three groups (NCI, MCI, AD) for brain samples and logistic regression across two groups (NCI, MCI/AD) for serum samples. The relationships between log-transformed brain metabolites levels with neurofibrillary tangle burden and neuritic plaque burden were expressed as boxplots across Braak scores (Kruskal-Wallis test) and CERAD scores (Wilcoxon rank-sum test), respectively. Using Spearman’s rank correlation test, we further evaluated the associations between the abundances of each identified metabolite and the global cognitive function score in both brain and serum samples. Linear regression models with each individual metabolite used as the predictor and each cognitive test as the response variable (adjusted for age, sex, years of education and presence of APOE ε4) were used to test the associations between metabolite and cognitive function. Similar analyses with an additional adjustment of BMI were conducted for serum samples. The Wilcoxon rank-sum test was carried out to explore whether identified variables were differentially expressed between NCI (converters) vs. NCI (non-converters), and between NCI (converters) vs. MCI/AD in sera. Then, we built a random forest (RF) predictive model to differentiate NCI (non-converters) vs. MCI/AD using glycolithocholate (GLCA), deoxycholate/cholate (DCA/CA) ratio, petroselinic acid, linoleic acid, myristic acid, palmitic acid, palmitoleic acid, and age as the predictors.

To differentiate MCI/AD vs. NCI (non-converters), we randomly split the data into 70% (training set) and 30% (testing set) 100 times. Each time, we trained an RF model on the training set to differentiate the MCI/AD from NCI (non-converters) and evaluated it on the testing set using the area under the receiver operating characteristic curve (AUROC), sensitivity (SE), and specificity (SP). A final model was built on the whole NCI (non-converters) and MCI/AD data. To investigate the pre-clinical predictive potentials as well as to validate the classification performance of our model, we utilized this model on the whole baseline NCI group to identify those NCI (converters) from NCI (non-converters). The differences of RF scores between NCI (non-converters) vs. NCI (converters), between NCI (non-converters) vs. MCI/AD group were tested by the Wilcoxon rank-sum test. To determine whether RF scores could independently differentiate NCI (converters) from NCI (non-converters) in the presence of potential confounders, we used the logistic regression method with RF scores as the predictor adjusting for sex, years of education, APOE ε4, and BMI. We additionally fitted linear mixed effects models to evaluate correlations between RF scores with global cognitive function and each of the five cognitive domains separately with a random effects term for education level, BMI and fixed effects terms for RF score, sex, and APOE ε4.

For the personalized pathway level analyses, we extracted metabolite information from the Human Metabolome Database (HMDB)^44^ and metabolic pathway information from the Kyoto Encyclopedia of Genes and Genomes (KEGG) database^45^ to map affiliated metabolites to metabolic pathways. We used the *pathifier* algorithm^46^ to transfer metabolic level information of each sample to pathway level by generating a pathway dysregulation score (PDS). For each pathway, each sample was projected onto a directed principal curve^47^, which was yielded depending on leading principal components of the pathway, to optimally pass through the cloud of samples. PDS was the distance along the curve between the projection of each sample and that of NCI. Thus, PDS could capture the pathway-level extent of abnormality (increments or decrements) for each participant relative to those with NCI. We performed similar data analysis on pathway level data to what we did on metabolomics level data. We tried to identify differential pathways using ordinal logistic regression across NCI, MCI, and AD groups in brain samples and logistic regression for NCI and MCI/AD in serum samples. Next, we explored the associations between identified pathways with neuropathology (Kruskal-Wallis test for Braak scores, Wilcoxon rank-sum test for CERAD scores) and cognitive performance (Spearman’s rank correlation test for global cognitive function, linear regression with adjustments for each cognitive test). Then, we examined the predictive potential of identified pathways in serum samples using univariate analysis (Wilcoxon rank-sum test for NCI (converters) vs. NCI (non-converters), NCI (converters) vs. MCI/AD). Finally, we built RF models on 70% training sets and tested them on 30% testing sets according to 100 times random splitting on the model construction data, and applied the final model on the validation data using ROC, SE, SP as evaluation methods.

Data were analyzed using R version 3.5.1 with packages including *pROC, pathifier, randomForest, ggplot2, ggsignif*, and *MASS*. The statistically significance was determined by a threshold of p values < 0.05.

## Results

### Participants and characteristics

For the joint analyses of the ROS/MAP study, we measured metabolomics of 566 serum samples (446 NCI, 109 MCI, and 11 AD at the time of blood draw) and 109 postmortem brain tissues from the dorsolateral prefrontal cortex (51 NCI, 31 MCI, and 27 AD at the time of death). A total of 92 participants had both brain and blood metabolomics data. NCI participants (n=446) are further categorized into 90 “NCI (converters)” and 356 “NCI (non-converters)”. NCI (converters) were those participants who were NCI at the time of blood draw and then experienced cognitive decline (MCI or AD), while NCI (non-converters) were participants who remained cognitively normal over follow-up. Detailed demographic characteristics of the serum samples and postmortem brain samples are included in Table 1. Among participants with postmortem brain samples, AD patients were more likely to have least one APOE ε4 allele compared to the NCI group as expected. The mean age of NCI and MCI/AD group at the time of blood draw among serum samples was 80.77 years (SD: 7.37) and 86.30 (SD: 6.26), respectively. Similarly, the age and the percentage of APOE ε4 carriers were higher in the NCI (converters) group than the NCI (non-converters) group. We did not observe other significant demographic characteristics differences across clinical groups (Table 1).

**Table 1.** Detailed demographic characteristics of study samples.

| **Brain samples** |  |  |  |  |
| --- | --- | --- | --- | --- |
|  | **Overall** | **NCI** | **MCI** | **AD** |
| *N* | 109 | 51 | 31 | 27 |
| Age, mean (SD) | 90.40 (5.86) | 90.06 (5.56) | 90.72 (6.14) | 90.68 (6.28) |
| Male, *n* (%) | 31 (28.4) | 18 (35.3) | 9 (29.0) | 4 (14.8) |
| Education, mean (SD) | 15.08 (3.17) | 15.25 (3.60) | 14.65 (2.27) | 15.26 (3.23) |
| APOE ε4-carrier, *n* (%) | 22 (21.2) | 6 (12.2) | 7 (24.1) | 9 (34.6) * |
| **Serum samples** |  |  |  |  |
|  | **Overall** | **NCI** | **MCI/AD** |  |
| *N* | 566 | 446 | 120 |  |
| Age (years), mean (SD) | 81.94 (7.49) | 80.77 (7.37) | 86.30 (6.26) ** |  |
| Male, *n* (%) | 121 (21.4) | 93 (20.9) | 28 (23.3) |  |
| Education (years), mean (SD) | 15.71 (3.11) | 15.79 (3.12) | 15.42 (3.09) |  |
| APOE ε4-carrier, *n* (%) | 95 (19.9) | 68 (18.1) | 27 (26.5) |  |
| **Serum NCI samples** |  |  |  |  |
|  | **Overall** | **NCI (non-converters)** | **NCI (converters)** |  |
| *N* | 446 | 356 | 90 |  |
| Age (years), mean (SD) | 80.77 (7.37) | 79.73 (7.42) | 84.87 (5.54) *** |  |
| Male, *n* (%) | 93 (20.9) | 74 (20.8) | 19 (21.1) |  |
| Education (years), mean (SD) | 15.79 (3.12) | 15.87 (3.16) | 15.48 (2.94) |  |
| APOE ε4-carrier, *n* (%) | 68 (18.1) | 43 (14.8) | 25 (29.4) **** |  |

\* Chi-square test, $P$ -value $< 0.05$ comparing AD vs. NCI
\*\* Wilcoxon rank sum test, $P$ -value $< 0.05$ comparing MCI/AD vs. NCI
\*\*\* Wilcoxon rank sum test, $P$ -value $< 0.05$ comparing NCI (converters) vs. NCI (non-converters)
\*\*\*\* Chi-square test, $P$ -value $< 0.05$ comparing NCI (converters) vs. NCI (non-converters)
*Abbreviations:* NCI, no cognitive impairment; MCI, mild cognitive impairment; AD, Alzheimer's disease; APOE $\epsilon 4$ , apolipoprotein E epsilon 4 allele; SD, standard deviation.

### Identifying metabolites differentially expressed in participants with cognitive impairment

In this study, 177 metabolites and 164 metabolites (129 overlapping metabolites) were detected in brain tissues and serum samples, respectively (Table S7, Table S8). Amino acids, BAs, carbohydrates, organic acids, and fatty acids were the predominant types of annotated metabolites (accounted for 84.17% of all the metabolites in brain tissues, and 84.75% in serum samples) (Fig. 1a, 1b, left panel). A total of 7 metabolites (1 BA, 1 BA ratio, 1 organic acid known as a long-chain fatty acid, and 4 fatty acids) showed significant differences across clinical groups in both brain and serum samples (*P*-value < 0.05, ordinal logistic regression for brain samples, logistic regression for serum samples) (Fig. 1a, 1b, right panel). In brain tissues, increments of the levels of GLCA, DCA/CA ratio, petroselinic acid, linoleic acid, myristic acid, palmitic acid, and palmitoleic acid followed the pattern NCI<MCI<AD. We observed increments of GLCA and DCA/CA ratio and decrements of petroselinic acid, linoleic acid, myristic acid, palmitic acid, and palmitoleic acid in sera of MCI/AD compared to NCI (Table 2). Previous findings were further validated in 92 individuals who have both brain and serum samples. From NCI to MCI and AD groups, increments of identified metabolites were observed in brain samples (Table S11). The increasing trend of GLCA, DCA/CA ratio and decreasing trend of FFAs among MCI/AD group relative to the NCI group were detected in sera (Table S11).

**Table 2.** Levels of metabolites differentially expressed in participants with cognitive decline.

| **Brain samples** |  |  |  |
| --- | --- | --- | --- |
|  | **NCI** | **MCI** | **AD** |
| GLCA %, median<br>[IQR] | 0.44 [0.24,0.84] | 0.59 [0.25,1.22] | 0.86 [0.42,1.24]* |
| DCA/CA, median<br>[IQR] | 6.06 [2.57,11.79] | 9.14 [6.00,15.75] | 8.60 [4.26,31.96]* |
| Petroselinic acid,<br>median [IQR] | 1696.67<br>[1323.38,2166.37] | 1856.55<br>[1529.43,2374.30] | 2181.55<br>[1761.35,2617.07]** |
| Linoleic acid, median<br>[IQR] | 13.86 [10.64,18.15] | 14.37 [10.40,21.62] | 19.93 [15.08,29.43]*** |
| Myristic acid, median<br>[IQR] | 164.14<br>[130.79,205.57] | 177.10<br>[136.34,205.38] | 206.86<br>[177.15,246.65]** |
| Palmitic acid, median<br>[IQR] | 8504.53<br>[6898.86,10496.19] | 9025.76<br>[7811.72,10964.45] | 10323.59<br>[9204.13,12785.95]*** |
| Palmitoleic acid,<br>median [IQR] | 64.96 [48.13,79.72] | 75.39<br>[61.45,100.90] | 82.06 [64.21,127.76]** |
| **Serum samples** |  |  |  |
|  | **NCI** | **MCI/AD** |  |
| GLCA %, median<br>[IQR] | 0.96 [0.49, 1.60] | 1.10 [0.54, 1.94] |  |
| DCA/CA, median<br>[IQR] | 10.30 [3.11, 28.10] | 14.78 [3.69, 44.75]<sup>#</sup> |  |
| Petroselinic acid,<br>median [IQR] | 0.84 [0.45, 1.41] | 0.49 [0.33, 1.02]<sup>###</sup> |  |
| Linoleic acid, median<br>[IQR] | 0.90 [0.64, 1.24] | 0.73 [0.58, 0.99]<sup>##</sup> |  |
| Myristic acid, median<br>[IQR] | 0.84 [0.59, 1.22] | 0.63 [0.52, 0.97]<sup>###</sup> |  |
| Palmitic acid, median<br>[IQR] | 0.89 [0.68, 1.18] | 0.78 [0.62, 1.00]<sup>##</sup> |  |
| Palmitoleic acid,<br>median [IQR] | 0.75 [0.35, 1.35] | 0.43 [0.23, 0.87]<sup>###</sup> |  |

\* $P$ -value < 0.05, \*\* $P$ -value < 0.01, \*\*\* $P$ -value < 0.001, by Wilcoxon rank sum test comparing AD vs. NCI

**Figure 1.**
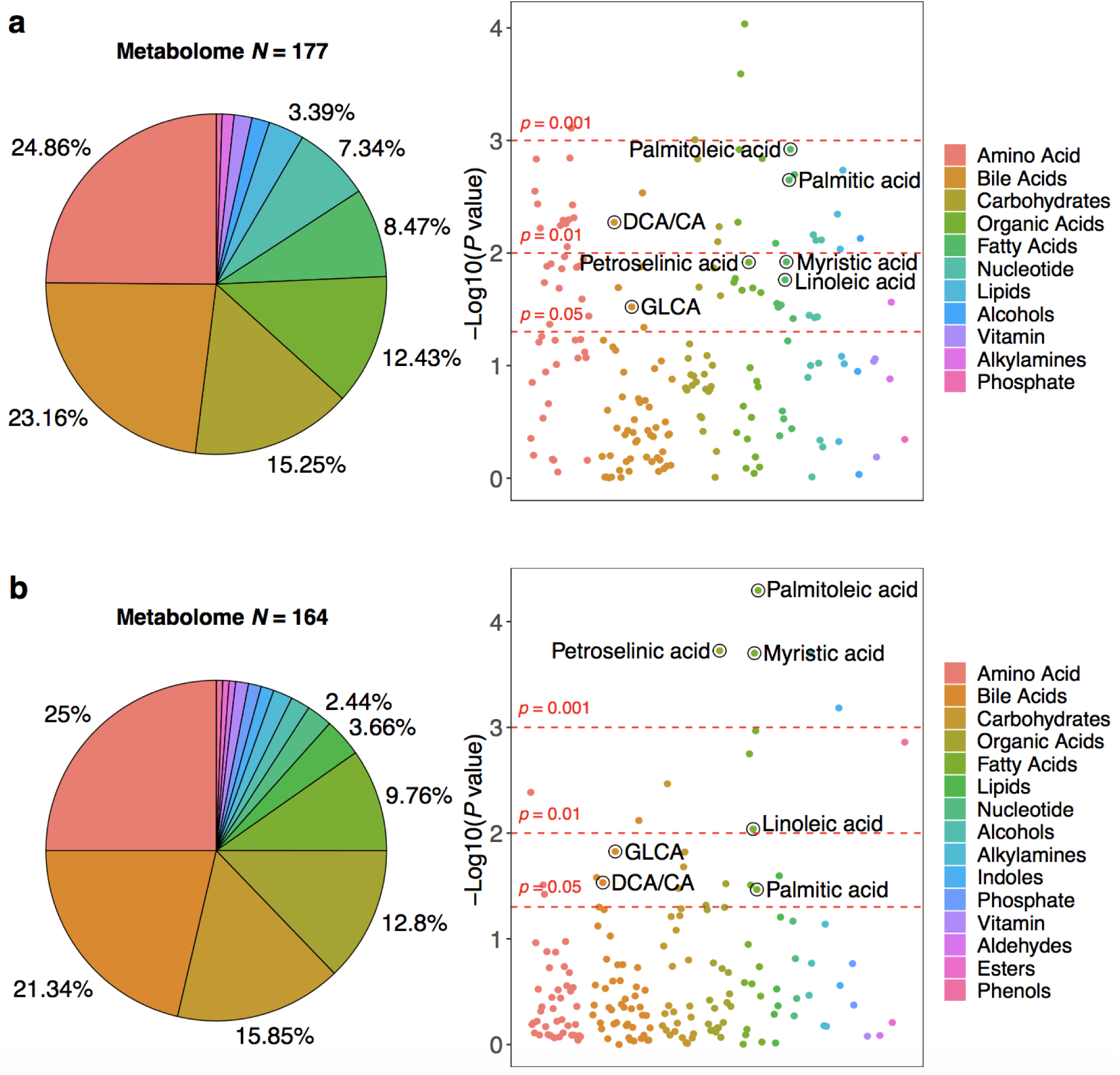
Brain metabolome and serum metabolome composition and alterations. (a) Left panel: the brain metabolome composition. Right panel: –log10(*P*-value) across clinical groups of brain tissues (NCI, MCI, AD). (b) Left panel: the serum metabolome composition. Right panel: –log10(*P*-value) across clinical groups of serum tissues (NCI, MCI/AD).

The seven brain metabolites were all negatively correlated with global cognitive function where higher scores indicate better cognitive performance (ρ = −0.091 for GLCA; ρ = −0.21 for DCA/CA ratio, ρ = −0.16 for petroselinic acid, ρ = −0.25 for linoleic acid, ρ = −0.22 for myristic acid, ρ = −0.2 for palmitic acid, and ρ = −0.26 for palmitoleic acid) using Spearman’s rank correlation analysis (Fig. 2a). Similarly, after adjusting for age, sex, years of education and APOE ε4, all identified metabolites remained negatively correlated with tests in five cognitive domains and the MMSE (see Table S2 for significant correlation pairs). The serum concentration of two BAs showed negative correlations with global cognitive function (ρ = −0.074 for GLCA; ρ = −0.075 for DCA/CA ratio), conversely fatty acids demonstrated positive correlations (ρ = 0.17 for petroselinic acid, ρ = 0.13 for linoleic acid, ρ = 0.13 for myristic acid, ρ = 0.11 for palmitic acid, and ρ = 0.17 for palmitoleic acid) (Fig. 2b). Linear regression revealed similar, consistent results in serum samples with adjustment for age, sex, years of education, APOE ε4, and BMI (see Table S2). Results of associations between identified metabolites/ratio and each cognitive domain are shown in Table S13. Correlations with global cognitive function were further validated in 92 individuals with both brain and serum samples and the directions were consistent with our previous findings among the all subjects. Seven identified metabolites were all negatively correlated with global cognitive function in brain samples, while two BAs showed negative correlations and five FFAs showed positive correlations in serum samples (Fig. S6). Additionally, the serum/brain ratio of identified FFAs were positively correlated with global cognitive test (a.k.a., lower levels of identified FFAs in serum and higher levels of identified FFAs in brain were associated with worse cognition) (Fig. S7).

**Figure 2.**
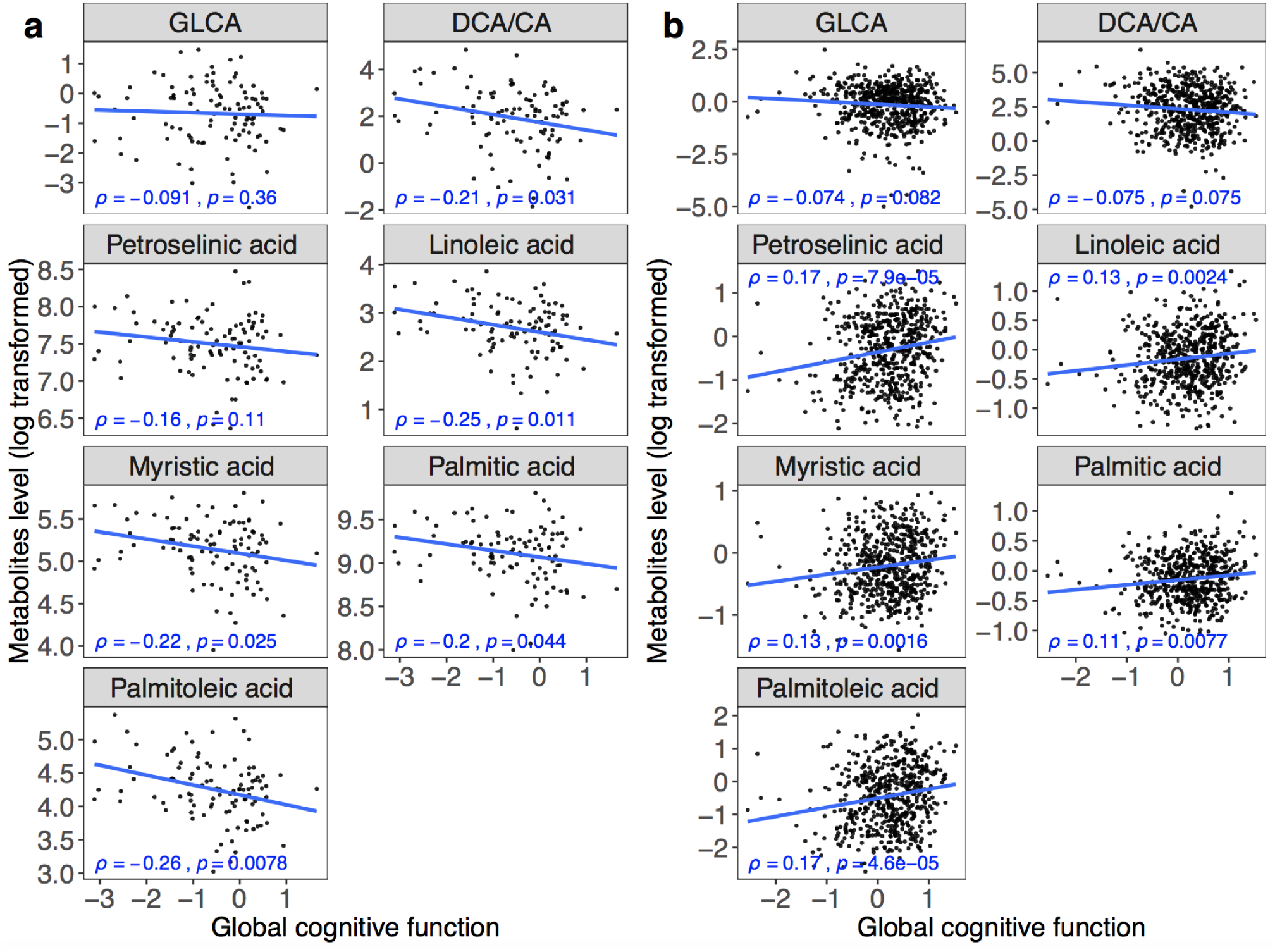
Associations between metabolites level and global cognitive function. (a) Boxplots showing group differences and *P* values for identified metabolites across Braak groups for brain tissue abundances. (b) Boxplots showing group differences and significances for identified metabolites across CERAD groups for brain tissue abundances. ρ, correlation coefficient of Spearman’s rank correlation test.

### Identified metabolites predicted antecedent cognitive impairment before the manifestation of clinical symptoms

The concentrations of GLCA and DCA/CA were significantly lower in the NCI (non-converters) group than in the NCI (converters) group. By contrast, the abundances of petroselinic acid, linoleic acid, myristic acid, palmitic acid, and palmitoleic acid were higher in the NCI (non-converters) group than in the NCI (converters) group (Fig. 3a). There were no significant differences in these metabolites between participants in NCI (converters) group vs. MCI/AD group (Fig. 3a). Using the seven metabolites and age, we built RF models on the 70% training set according to 100-times randomly splitting approach to differentiate MCI/AD patients from NCI (non-converters) group. The median of 100 times AUC on 30% testing set was 0.772 (95% CIs = 0.763-0.781) with 0.786 SE (95% CIs = 0.767-0.804) and 0.716 SP (95% CIs = 0.695-0.737) using Youden’s index to maximize the sum of SE and SP (Fig. 3b). RF models showed decent classification performances in differentiating MCI/AD group from NCI (non-converters). Next, we were interested in studying the model’s early diagnostic capability for predicting NCI (converters) before clinical diagnosis. The model was thus applied on the whole NCI group at baseline to differentiate NCI (converters) from NCI (non-converters). We achieved an AUC of 0.804 (95% CIs = 0.749-0.859, SE = 0.722, SP = 0.749 at the cutoff value of 0.357) (Fig. 3c) with significant differences in RF scores between NCI (converters) vs. NCI (non-converters), between NCI (non-converters) vs. MCI/AD group using the Wilcoxon rank-sum test (*P*-value < 0.001) (Fig. 3d). After further adjustment for sex, years of education, APOE ε4, and BMI, the RF scores remained significant (as an independent predictor) with a coefficient of 6.994 (*P* value < 0.001) (Table S3). Additionally, the RF scores showed significant negative correlations with global cognitive function and five cognitive domains with same adjustment in mixed effects models (Table S12).

**Figure 3.**
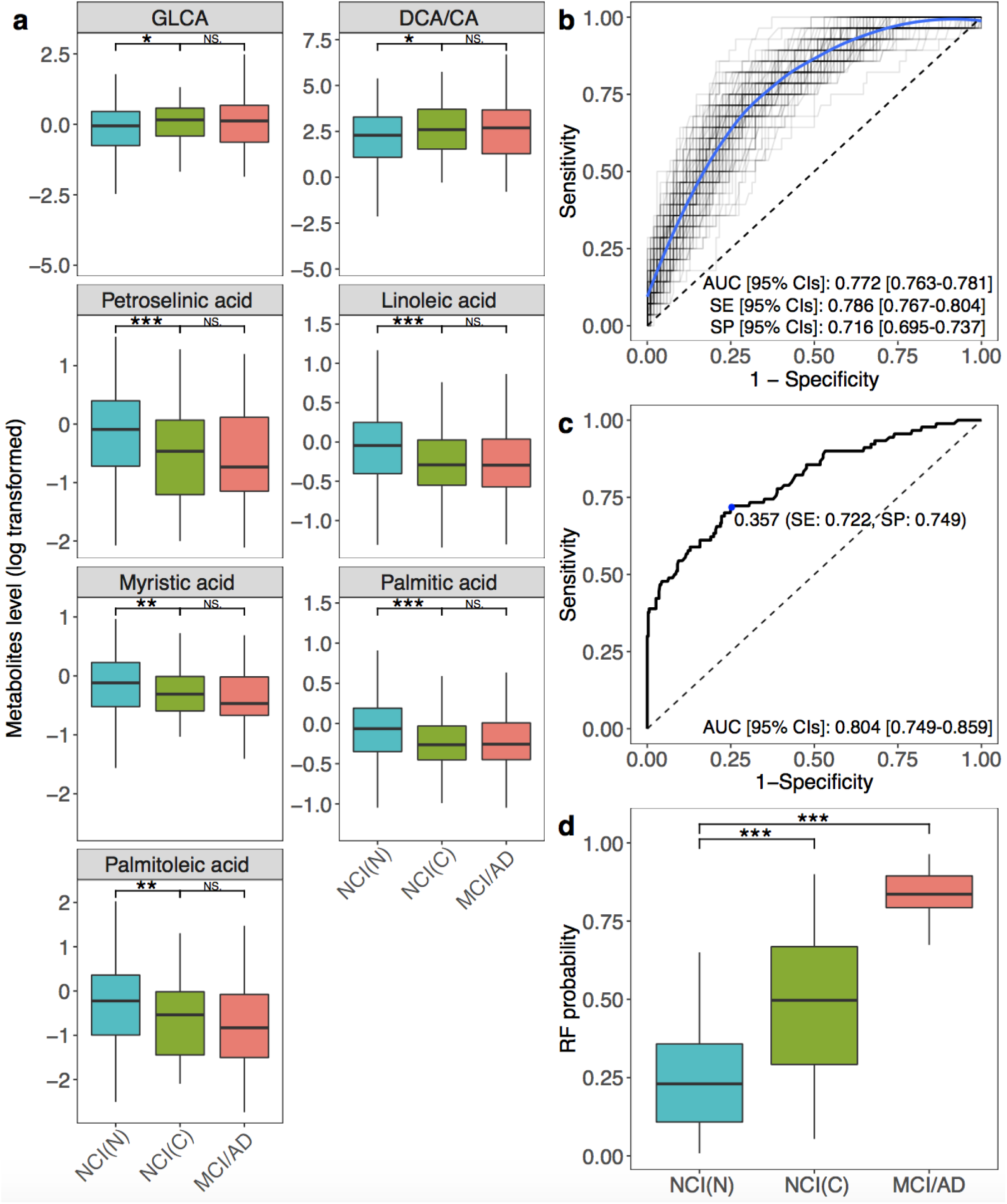
The identified panel of metabolites and its predictive performance. (a) Boxplots showing group differences and *P* values for identified metabolites across NCI (non-converters), NCI (converters), and MCI/AD for serum abundances. (b) ROC curves of metabolite models trained on the 70% training data and tested on the 30% testing data according to 100-times randomly training-testing splitting. (c) The ROC curve of the final metabolite model on the validation data. (d) RF scores of the final metabolite model across NCI (non-converters), NCI (converters), and MCI/AD. * *P*-value < 0.05, ** *P*-value < 0.01, *** *P*-value < 0.001, Wilcoxon rank sum test. The optimal cutoff was determined by the Youden index. *Abbreviations*: AD, Alzheimer’s disease; AUC, area under the receiver operating characteristic curve; NCI(C), NCI (converters); NCI(N), NCI (non-converters); CIs, confidence intervals; MCI, mild cognitive impairment; NS, not significant; SE, sensitivity; SP, specificity.

### Personalized metabolic pathway-based study for the association and prediction of cognitive impairment

Considering altered metabolite levels were significantly associated with cognitive impairment and showed early predictive value of clinical symptoms onset, we then employed the *pathifier* algorithm to summarize metabolite information to pathways level for further examinations. All PDS scores range from 0 to 1, where larger scores represent the higher extent of the abnormality in the corresponding metabolic pathway. A large number of metabolites (109/177 metabolites in brain tissues and 102/164 metabolites in sera) were successfully mapped to the KEGG metabolic pathways. This method identified 52 metabolic pathways in brain tissues and 45 metabolic pathways in serum samples (44 overlapping pathways) (Fig. S1a, S1b, left panel; Table S9, Table S10), three of which (i.e., primary BAs biosynthesis, FFAs biosynthesis, and biosynthesis of unsaturated FFAs) were significantly shifted in both brain and serum samples (*P*-value < 0.05, ordinal logistic regression for brain samples, logistic regression for serum samples) (Fig. S1a, S1b, right panel). We noted increased PDS for all three identified pathways from NCI to MCI/AD that suggested dysregulation of these metabolic pathways in MCI/AD patients compare to NCI. Detailed PDS of these pathways stratified by diagnostic groups are described in Table S4. Results also indicated that higher PDS were significantly associated with lower global cognitive function (i.e., worse cognitive performance) in both brain (ρ = −0.16 for primary BAs biosynthesis pathway, ρ = −0.23 for FFAs biosynthesis pathway, ρ = −0.26 for biosynthesis of unsaturated FFAs pathway) (Fig. S4) and serum samples (ρ = −0.16 for primary BAs biosynthesis pathway, ρ = −0.13 for FFAs biosynthesis pathway, ρ = −0.14 for biosynthesis of unsaturated FFAs pathway) (Fig. S5) respectively. In Table S5, we show the significant negative associations between each cognitive test and PDS of 3 pathways after adjusting for age, sex, years of education, and APOE ε4 (further adjustment for BMI in serum samples). Two fatty acid pathways showed significantly different PDS between the NCI (non-converters) group and the NCI (converters) group (*P*-value = 0.0012, and *P*-value < 0.001), respectively. A gradually increasing trend was noted for the BAs pathway across groups (i.e., NCI (non-converters) < NCI (converters) and MCI/AD) (Fig. 4a).

**Figure 4.**
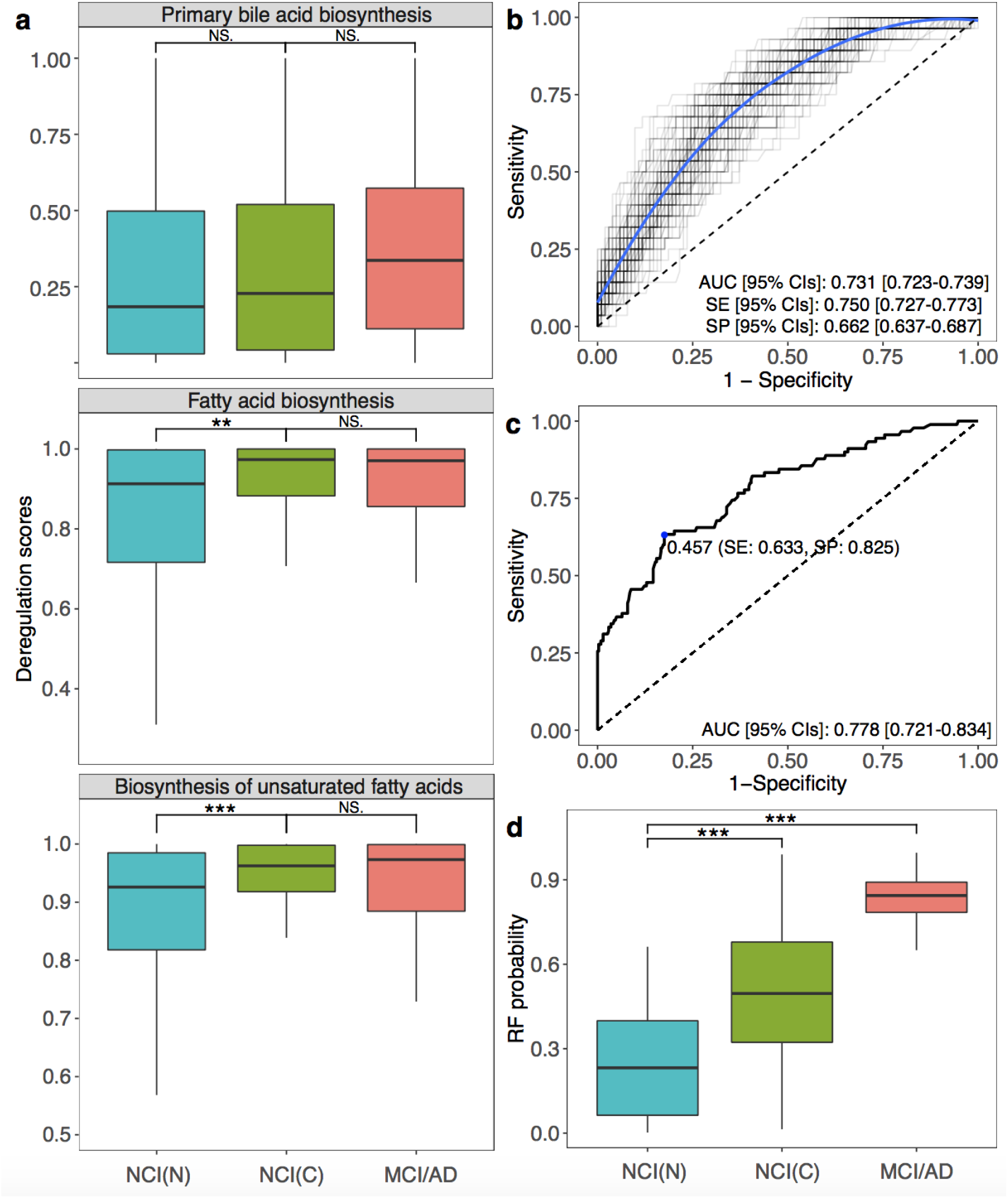
The pathway panel and its predictive performance. (a) Boxplots showing group differences and *P* values for identified pathways across NCI (non-converters), NCI (converters), and MCI/AD for serum abundances. (b) ROC curves of pathway models trained on the 70% training data and tested on the 30% testing data according to 100-times randomly training-testing splitting. (c) The ROC curve of the final pathway model on the validation data. (d) RF scores of the final pathway model across NCI (non-converters), NCI (converters), and MCI/AD. * *P*-value < 0.05, ** *P*-value < 0.01, *** *P*-value < 0.001, Wilcoxon rank sum test. The optimal cutoff was determined by the Youden index. *Abbreviations*: AD, Alzheimer’s disease; AUC, area under the receiver operating characteristic curve; NCI(C), NCI (converters); NCI(N), NCI (non-converters); CIs, confidence intervals; MCI, mild cognitive impairment; NS, not significant; SE, sensitivity; SP, specificity.

We then constructed a discriminant RF model in 70% training data and tested on 30 testing data based on three identified metabolic pathways along with age to differentiate MCI/AD from NCI (non-converters) in model construction data using 100-times randomly splitting approach. The median AUC on testing set was 0.731 (95% CIs = 0.723-0.739) with 0.750 SE (95% CIs = 0.727-0.773) and 0.662 SP (95% CIs = 0.637-0.687) (Fig. 4b). Applying the RF model to the whole NCI data at baseline could successfully discriminate NCI (converters) from NCI (non-converters) with an AUC of 0.778 (95% CIs = 0.721-0.834), SE = 0.633, SP = 0.825, cutoff value = 0.457 (Fig. 4c). Similarly, predictive RF scores were significantly different between NCI (converters) vs. NCI (non-converters), and NCI (non-converters) vs. MCI/AD group (*P*-value < 0.001) (Fig. 4d). After adjusting for sex, years of education, APOE ε4, and BMI, the RF scores remained statistically significant as an independent predictor with a coefficient of 5.294 (*P*-value < 0.001) (Table S6).

### Identified metabolites and metabolic pathway were associated with neuropathology

Notably, we observed significant differences (gradually increasing trend) of metabolite abundances in brain tissue across five Braak score groups: DCA/CA ratio (*P*-value = 0.021), petroselinic acid (*P*-value = 0.038), linoleic acid (*P*-value = 0.041), palmitic acid (*P*-value = 0.043), and palmitoleic acid (*P*-value = 0.038) (Fig. S8a). Consistently, the abundance of six metabolites in the brain were higher in the CERAD AD group (score 1∼2) than the CERAD non-AD group (score 3∼4): GLCA (*P*-value = 0.041), petroselinic acid (*P*-value = 0.014), linoleic acid (*P*-value < 0.001), myristic acid (*P*-value = 0.015), palmitic acid (*P*-value = 0.026), and palmitoleic acid (*P*-value = 0.0019) (Fig. S8b). Brain PDS of identified pathways were gradually increased across five Braak score groups, where the biosynthesis of unsaturated FFAs pathway differed with statistical significance (*P*-value = 0.036) (Fig. S2). Higher brain PDS were associated with a significantly greater risk of neuritic plaque burden based on CERAD criteria: primary BAs biosynthesis (*P*-value = 0.04), FFAs biosynthesis (*P*-value = 0.028), and biosynthesis of unsaturated FFAs (*P*-value = 0.0089) (Fig. S3).

## Discussion

The alternations of metabolic profiles, especially in the brain of MCI and AD patients have not been extensively studied. In our study, we comprehensively measured 164 and 177 metabolites in serum and brain samples, respectively. Our findings showed six significantly differential metabolites and one ratio in both brain and serum samples (petroselinic acid, linoleic acid, myristic acid, palmitic acid, and palmitoleic acid, GLCA, and DCA/CA ratio). The identified metabolites/ratio showed consistent associations with neuropathology results, and cognitive tests regardless of adjustments for potential confounding factors (i.e., age, sex, years of education, APOE ε4, and BMI). Additionally, these metabolites demonstrated group differences of NCI (converter) vs. NCI (non-converter) within baseline normal controls which indicated their potential predictive values. A personalized pathway-level analysis further demonstrated our findings of FFAs and BAs metabolic perturbations among cognitively impaired patients and the predictive values of using personalized pathway information.

Altered FAs profiles were found in different regions of brain tissue in AD patients and linked to neuropathology and cognitive performance^14^. Our study showed five significant increments of brain FFAs, while decrements of these FFAs appeared in the serum of AD patients. The serum/brain ratio of identified FFAs were positively correlated with global cognitive test (i.e., lower levels of identified FFAs in serum and higher levels of identified FFAs in brain were associated with worse cognition) (Fig. S7).

Our findings are consistent with prior observations in AD mice that the significant accumulation of FFAs in the hippocampus and cortex, including palmitoleic acid, palmitic acid, and linoleic acid, might be associated with the utilization of FFAs in brain^25^. It is well-known that brain is one of the most energy-demanding organs with a high glycolytic catabolizing rate of glucose consumption. The energy supply shift from glucose towards alternative energy sources (e.g., ketone bodies and FFAs) has been observed in other neuropsychiatric conditions^48^ (Fig. 5). More interestingly, in the early stage of AD reduced glucose utilization and metabolic dysfunction can be detected using FDG-PET, which is one of the earliest detectable symptoms of the disease^49^. The metabolic instability with decreased glucose utilization in impaired neurons among AD patients occurs up to twenty years prior to the onset of clinical symptoms indicating that metabolic decline may contribute to the development of cognitive impairment^50^. Previous studies suggested that fatty acid can across the blood-brain barrier (BBB) via simple diffusion. At the same time, fatty acid transport proteins are also involved in this process. Fatty acid binding protein 5 (FABP-5), fatty acid transport proteins-1 (FATP-1), FATP-4 and fatty acid translocase (CD36) are the key FFA transport proteins and these transporters are expressed in human brain microvessel endothelial cells (HBMEC)^51^. The significant decrement of the movement for many FFAs (including linoleic acid, myristic acid, and palmitic acid) from apical medium to the basolateral medium across HBMEC monolayer with the knockdown of these FFA transport proteins was observed^51^. Additionally, many studies demonstrated that the CD36 gene is associated with AD^52,53^, and the BBB damage may also facilitate the transferring of FFAs from blood to brain in AD patients^54^.

**Figure 5.**
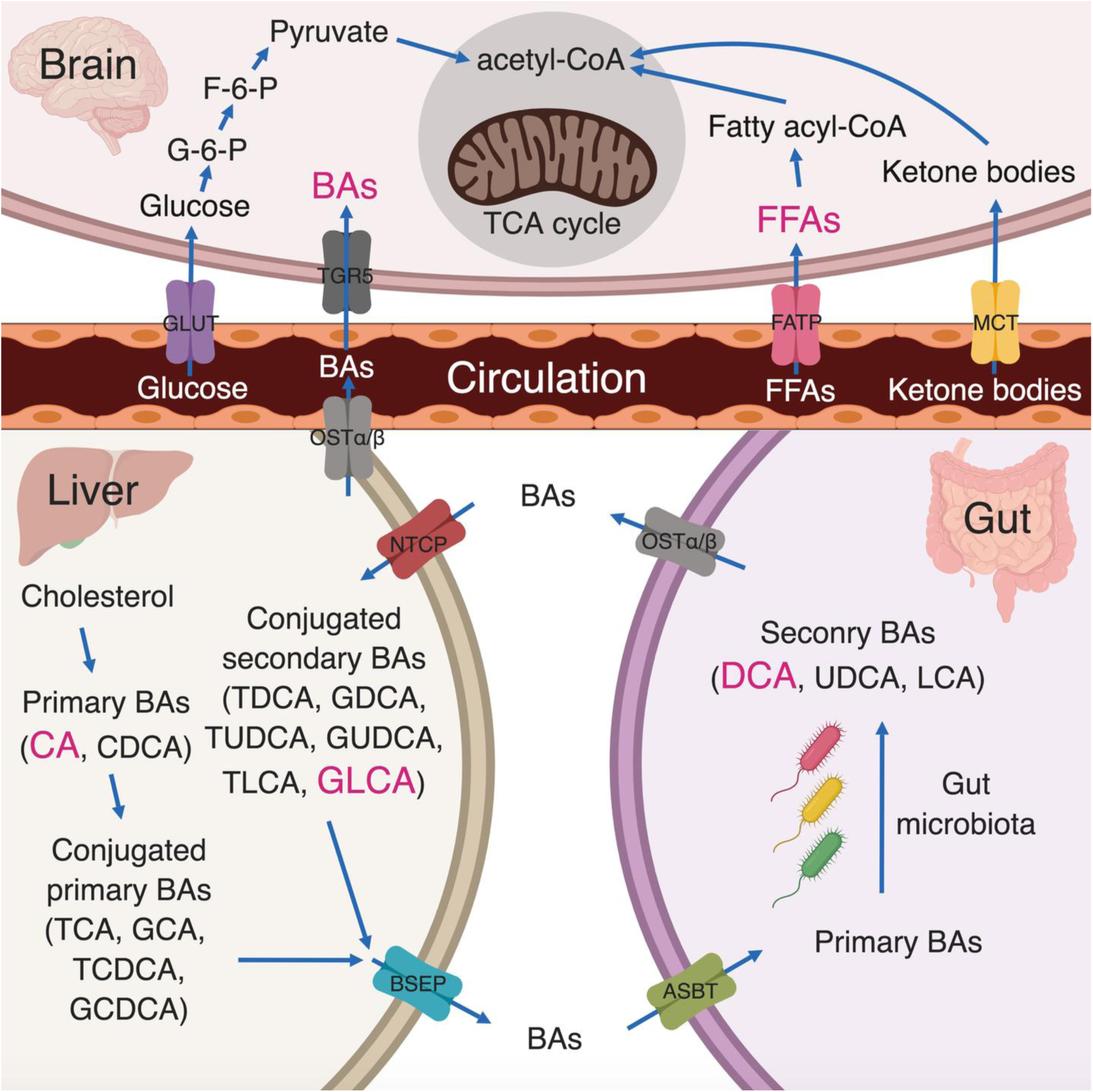
Pathways involved in FFAs and BAs. Healthy neurons are highly glycolytic, catabolizing rate of glucose consumption through the glycolysis and tricarboxylic acid cycle to produce ATP. Reduced glucose utilization and metabolic dysfunction could be detected using FDG-PET and metabolomics approaches in AD patients. The metabolic instability with decreased glucose utilization in impaired neurons can cause the energy supply shift towards alternative energy sources, e.g., FFAs and ketone bodies. Primary BAs are synthesized in the liver from cholesterol. A dysfunction of gut microbiome can cause the accumulation of cytotoxic secondary bile acids, e.g., DCA and GLCA, which can be secreted into the systemic circulation and then across the blood-brain barrier to enter the brain. *Abbreviations*: ASBT, apical sodium-dependent bile acid transporter; BSEP, bile salt export pump; CA, cholate; CDCA, chenodeoxycholate; CoA, coenzyme A; DCA, deoxycholate; F-6-P, fructose-6-phosphate; FATP, fatty acid transporter; G-6-P, glucose-6-phosphate; GCA, glycocholate; GCDCA, glycochenodeoxycholate; GDCA, glycodeoxycholate; GLCA, glycolithocholate; GLUT, glucose transporter; GUDCA, glycoursodeoxycholate; LCA, lithocholate; MCT, monocarboxylate transporter; NTCP, sodium/taurocholate co-transporting polypeptide; OST, organic solute and steroid transporter; TCA, taurocholate; TCDCA, taurochenodeoxycholate; TDCA, taurodeoxycholate; TLCA, taurolithocholate; TUDCA, tauroursodeoxycholate; TGR5, G protein–coupled bile acid receptor; UDCA, ursodeoxycholate.

In addition to the FFAs dysregulation, the association between cholesterol metabolism and AD leads to substantial research interests of how BAs profile changes in cognitively impaired individuals and AD patients^55^. Furthermore, BAs play a major role in regulating energy homeostasis through binding to nuclear receptors and both primary and secondary BAs can across the BBB via a gut-liver-brain axis^56,57^. In this study, we observed one secondary conjugated BA (i.e., GLCA) and one primary-secondary BAs ratio (DCA/CA) increased in both serum and brain tissue in MCI/AD patients compared to NCI, which was consistent with previous findings^12^. Primary BAs are synthesized in the liver from cholesterol and are then bio-transformed into secondary BAs by the gut microbiota (Fig. 5). The observed increasing DCA/CA ratio here might suggest a dysfunction of bacterial 7α-dehydroxylases which leads to the accumulation of cytotoxic secondary bile acids. Similarly, GLCA, another cytotoxic BA, was increased in our study. The survival analysis conducted by MahmoudianDehkordi *et al*. on MCI conversion to AD showed significantly changed prognostic endpoints when splitting samples into different groups according to BA levels^12^, which also suggested that the total BAs/BA ratios could serve as potential predictors for the conversion of ADs among non-AD clinical groups.

This study has limitations. First, the sample size of brain tissues in this study is relatively limited due to the study design which required available ante-mortem MRI. Additional work with a larger number of brain tissues is ongoing. Second, the current serum metabolomics data is cross-sectional therefore precluding explorations of changes in the serum metabolome changes over time. Third, although we validated our prediction models on the whole NCI group to show its additional predictive values in differentiating NCI (converters) from NCI (non-converters), more external validation data are needed to validate our results. Fourth, our findings suggest the emerging role for gut microbiome in the development of AD. Thus, further experimental microbiota studies could provide a better understand of how the gut-brain axis plays a role in AD development. Last, since our results provide a proof-of-concept for the feasibility of early detection among NCI subjects at high risk of developing cognitive impairment, future clinical studies should be designed to explore the benefits of early interventions. The study also has strengths in that the follow-up and autopsy rates in the parent cohorts are very high leading to excellent internal validity. The diagnostic groups were comparable as they came from a single larger cohort.

Collectively, our findings present a new point of view into the pre-clinical evolution of AD and lend strength to the hypothesis that individuals with higher risks of cognitive impairment can be identified before the development of overt symptoms via a metabolomics approach. To this end, we provide two predictive models: one based on differentially expressed metabolites and the other on identified metabolic pathways. Varma *et al*. utilized machine learning approaches to identify potential metabolites related to AD pathology and progression^17^. We took their work one step further by constructing machine learning models to discriminant the final-state healthy controls vs. MCI/AD patients. The 100 times resampling results demonstrated the feasibility and robustness of our predictive models. Particularly, by employing our models on the baseline group, we successfully identified the high-risk subgroup (i.e., NCI (converters)) several years before the clinical diagnosis.

## Acknowledgments

The results published here are in whole or in part based on data obtained from the AMP-AD Knowledge Portal (doi:10.7303/syn2580853). Study data provided by the Rush Alzheimer’s Disease Center, Rush University Medical Center, Chicago were supported through funding by NIA grants (P30AG10161, R01AG15819, R01AG17917, R01AG30146, R01AG36836, U01AG32984, U01AG46152), the Illinois Department of Public Health, and the Translational Genomics Research Institute. The NIA supported the AD Metabolomics Consortium which is a part of NIA’s national initiatives AMP-AD and M2OVE-AD (R01 AG046171, RF1 AG051550, and 3U01AG024904-09S4). RKD and MA are supported by NIA grants RF1 AG058942 and R01 AG057452. The study was supported by NIA grants U01AG46152.

## Author contributions

W.J. led the study along with RKD lead PI of Alzheimer Disease Metabolomics consortium. DB, De Jager, RKD designed study and enabled linking of biochemical data to clinical data. J.W., R.W., SMD and G.X. performed the data analysis, and implemented the methodology. J.W and R.W. prepared the original manuscript. G.X. performed the metabolomics analysis. Data management and medication term mapping were carried out by C.B.. Biochemical interpretation was done by M.A., R.B., X.H., R.K.D., W.J. Technical and bibliographic research was provided by G.L. J.W, R.W., G.X., W.J. M.A., G.L. D.B., P.L.D. and R.K.D. reviewed and edited the final manuscript.

