## Supplemental Figures for "Peripheral Serum Metabolomic Profiles Inform Antecedent Central Cognitive Impairment in Older Adults"

**Checklist of Supporting Information**

**Figure S1.** Brain metabolic pathways and serum metabolic pathways composition and alterations

**Figure S2.** PDS of metabolic pathways across Braak scores in brain

**Figure S3.** PDS of metabolic pathways across CERAD scores in brain

**Figure S4.** Correlations between PDS of metabolic pathways and global cognitive function in brain

**Figure S5.** Correlations between PDS of metabolic pathways and global cognitive function in sera

**Figure S6.** Associations between metabolites level and global cognitive function among participants with both brain and serum samples.

**Figure S7.** Associations between level of serum/brain FFAs ratios and global cognitive function among participants with both brain and serum samples.

**Figure S8.** Associations between metabolites level and Braak scores, CERAD scores.

**Supplementary Text.**

**Table S1.** List of cognitive performance tests

**Table S2.** Associations between identified metabolites/ratio and cognitive performance tests adjusting for age, gender, years of education, APOE ε4, BMI

**Table S3.** Logistic regression of metabolite marker panel-based RF score to discriminate NCI (converters) vs. NCI (non-converters) adjusting for gender, years of education, APOE ε4, and BMI

**Table S4.** PDS of metabolic pathways differentially expressed in participants with cognitive decline

**Table S5.** Associations between identified metabolic pathways and cognitive performance tests adjusting for age, gender, years of education, APOE ε4, and BMI

**Table S6.** Logistic regression of metabolic pathway panel-based RF score to discriminate NCI (converters) vs. NCI (non-converters) adjusting for gender, years of education, APOE ε4, and BMI

**Table S7.** Levels of detected metabolites across clinical groups in brain samples

**Table S8.** Levels of detected metabolites across clinical groups in serum samples

**Table S9.** PDS of mapped metabolic pathways across clinical groups in brain samples

**Table S10.** PDS of detected metabolic pathways across clinical groups in serum samples

**Table S11.** Levels of identified metabolites in samples with both brain and blood metabolomics data

**Table S12.** Mixed effects model of metabolite marker panel-based RF score adjusting for age, gender, years of education, APOE ε4, and BMI

**Table S13.** Associations between identified metabolites/ratio and cognitive performance domains adjusting for age, gender, years of education, APOE ε4, and BMI.

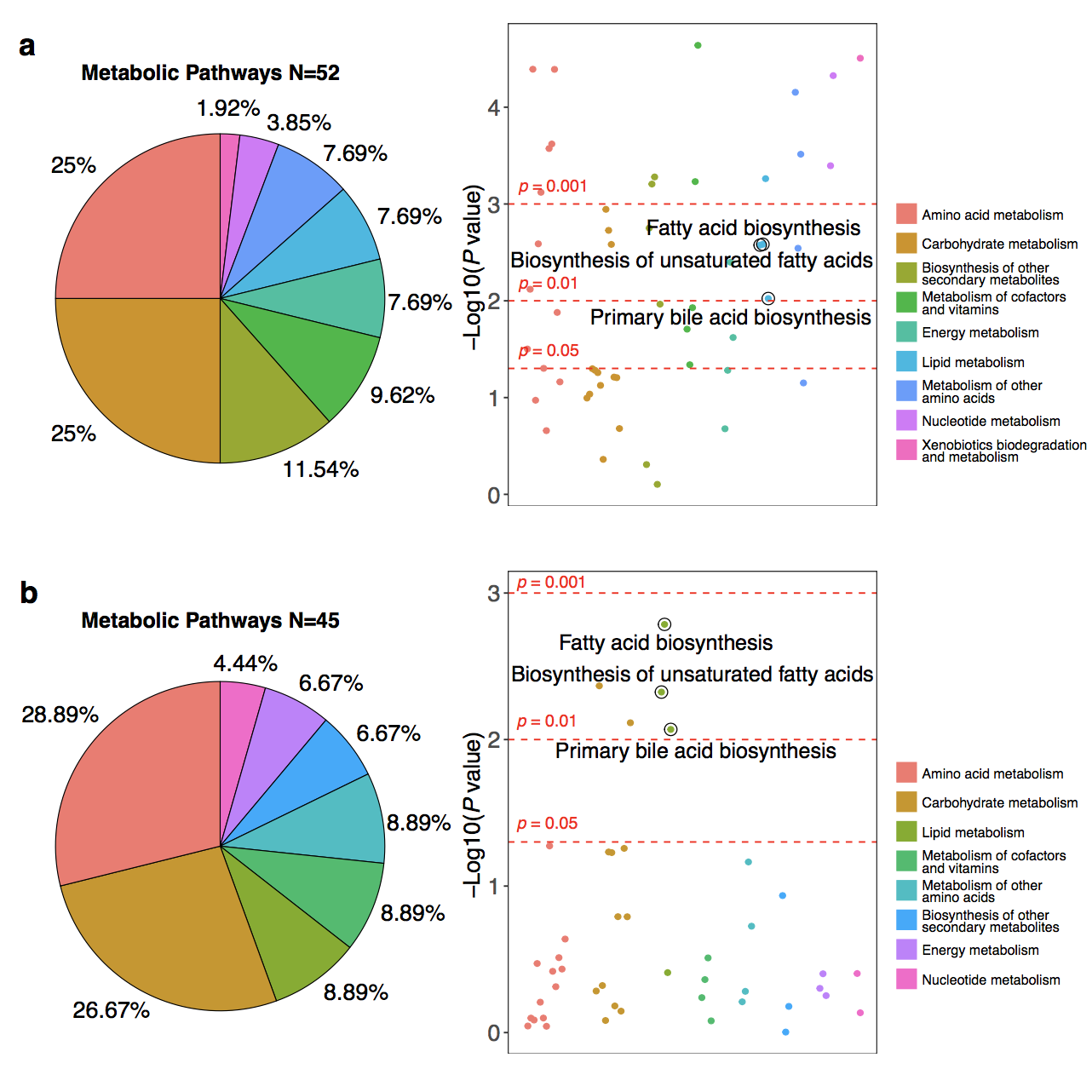

**Figure S1.** **Brain** **metabolic** **pathways and serum metabolic pathway compositions and alterations.**

(a) Left panel: the mapped pathway composition in brain. Right panel: –log10 (*P*-value) across clinical groups (NCI, MCI, AD). (b) Left panel: the mapped pathways composition in serum. Right panel: –log10 (*P*-value) across clinical groups (NCI, MCI/AD).

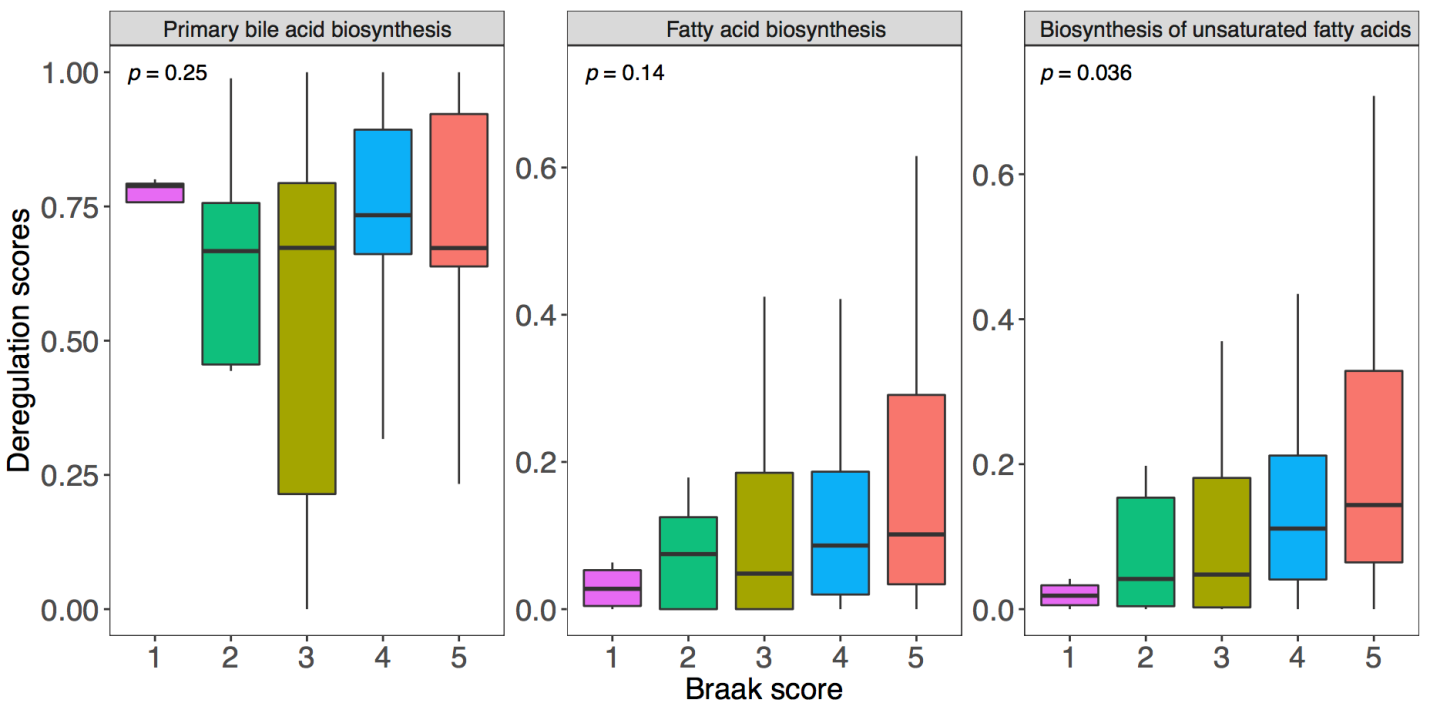

**Figure S2. PDS of metabolic pathways across** **Braak scores in brain.**

Boxplots showing group differences and *P*-values for identified pathways across Braak groups for brain tissues.

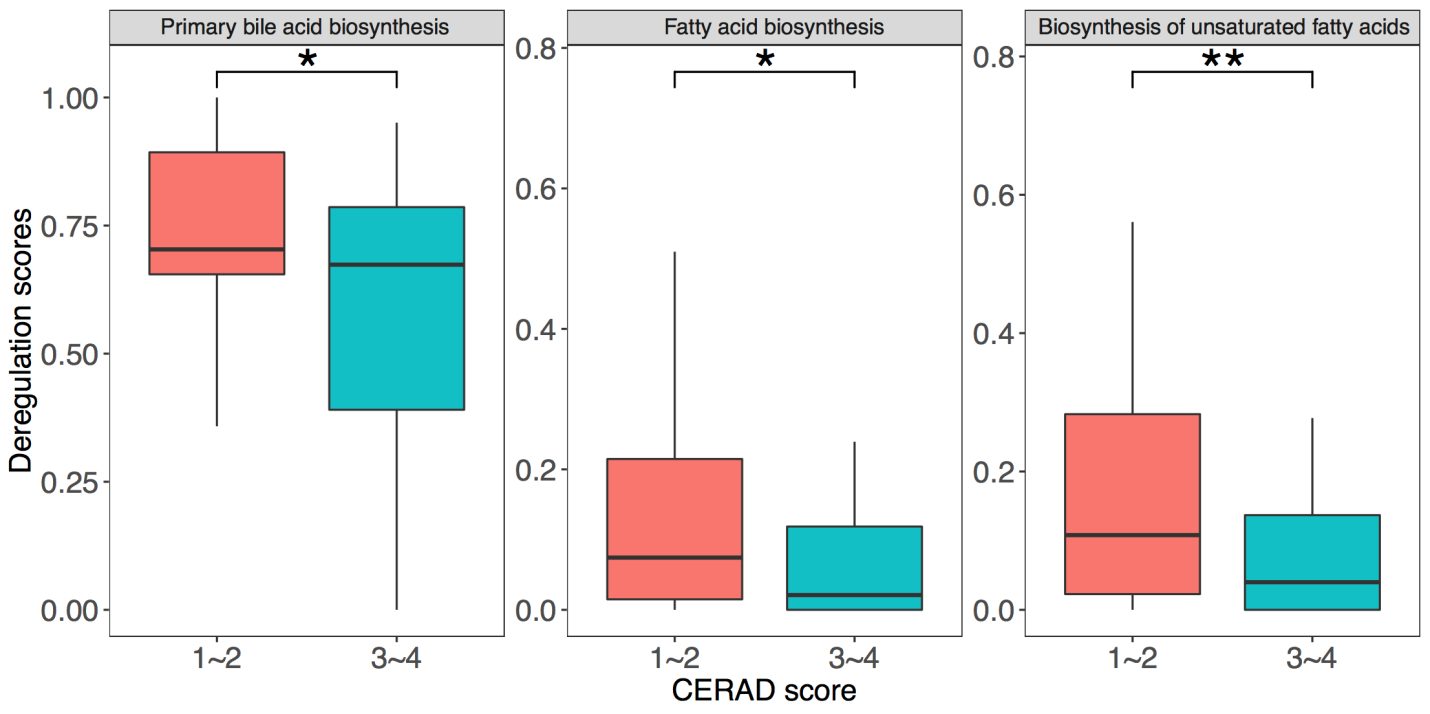

**Figure S3. PDS of metabolic pathways across** **CERAD scores in brain.**

Boxplots showing group differences and significances for identified pathways across CERAD groups for brain tissues. * *P*-value < 0.05, ** *P*-value < 0.01, *** *P*-value < 0.001, Wilcoxon rank sum test. CERAD, Consortium to Establish a Registry for Alzheimer’s Disease

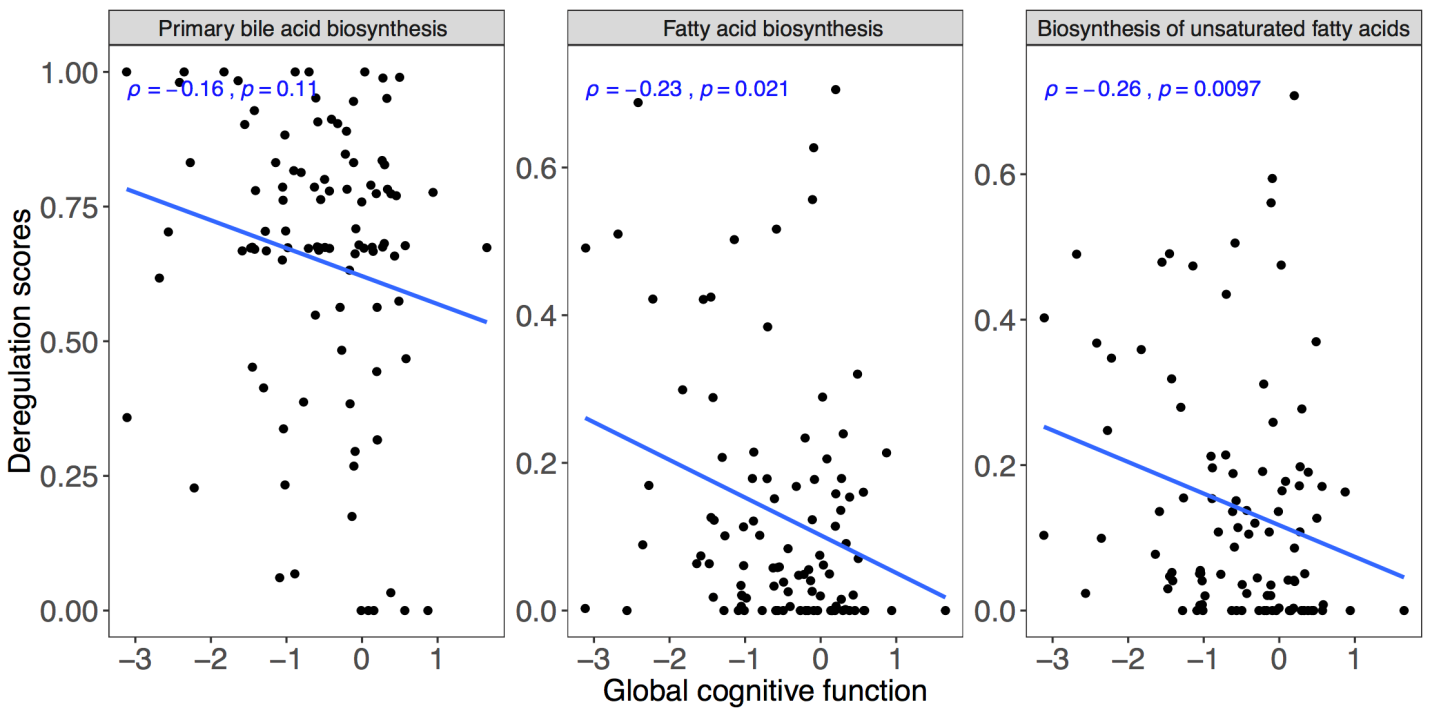

**Figure S4. Correlations between PDS of metabolic pathways and global cognitive function in brain.**

Scatterplots with ρs and *P*-values showing correlations between brain pathway PDS and global cognitive function. ρ, correlation coefficient of Spearman’s rank correlation test.

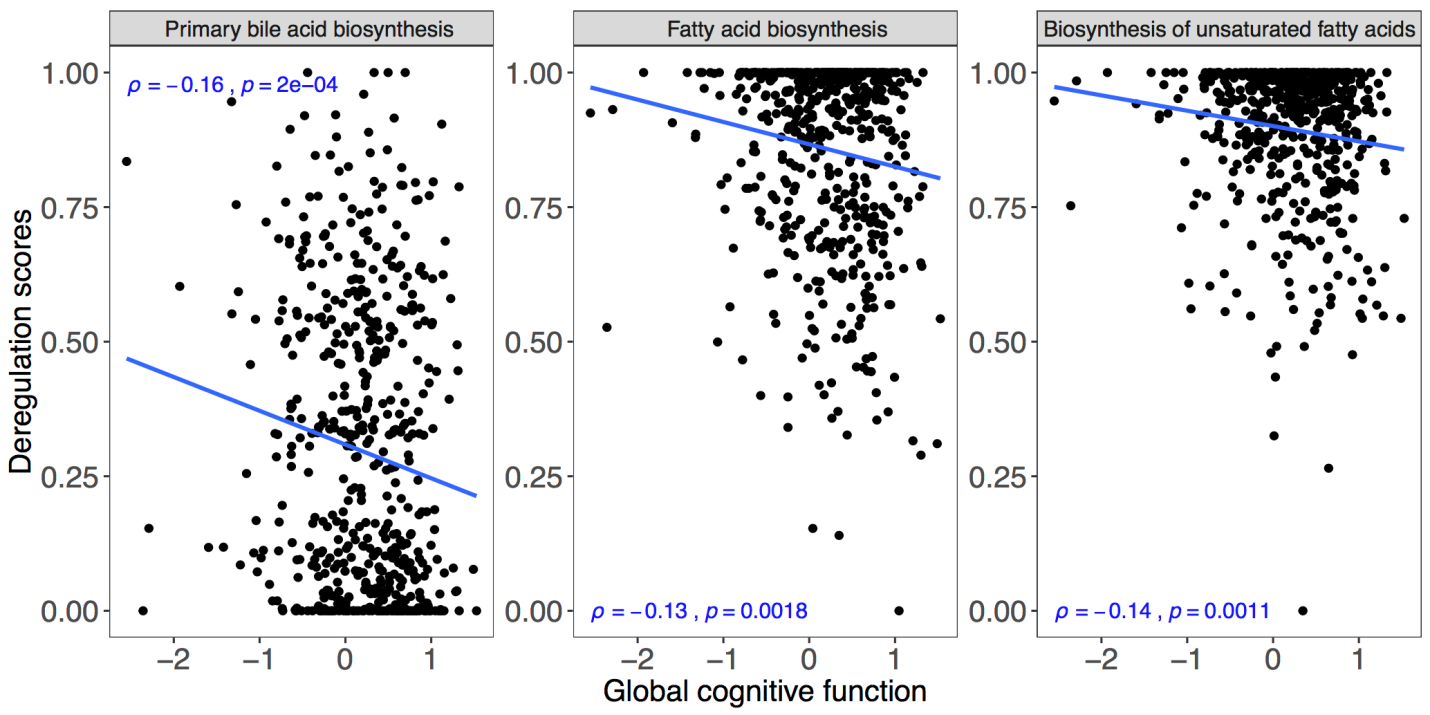

**Figure S5. Correlations between PDS of metabolic pathways and global cognitive function in sera.**

Scatterplots with ρs and *P*-values showing correlations between serum pathway PDS and global cognitive function. ρ, correlation coefficient of Spearman’s rank correlation test.

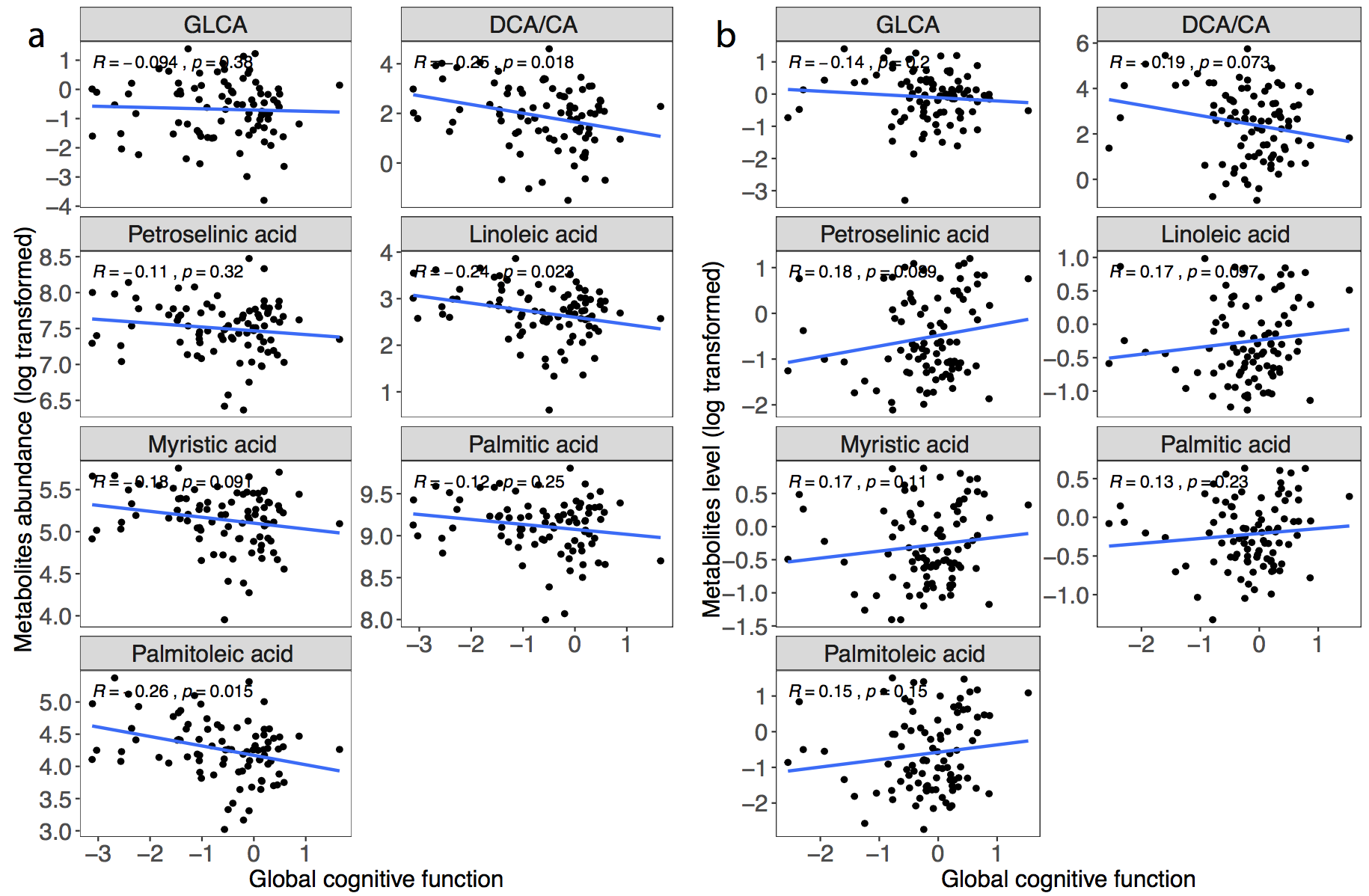

**Figure S6. Associations between metabolites level and global** **cognitive function among participants with both brain and serum samples.**

(a) Scatterplots with ρs and p-values showing associations between brain metabolites abundances and global cognitive function. (b) Scatterplots with ρs and p-values showing associations between serum metabolites abundances and global cognitive function. ρ, correlation coefficient of Spearman’s rank correlation test.

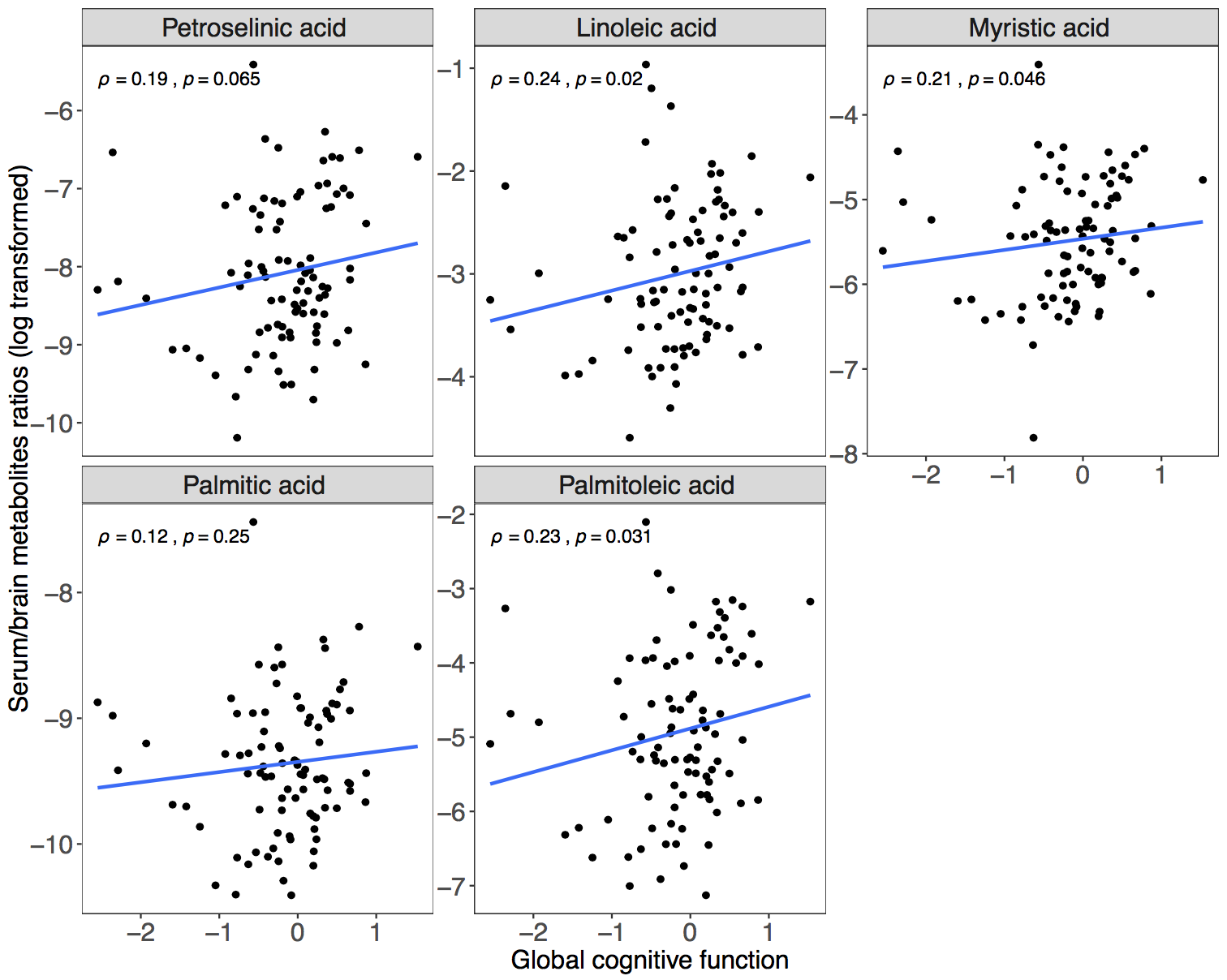

**Figure S7. Associations between level of serum/brain FFAs ratios and global cognitive function among participants with both brain and serum samples.**

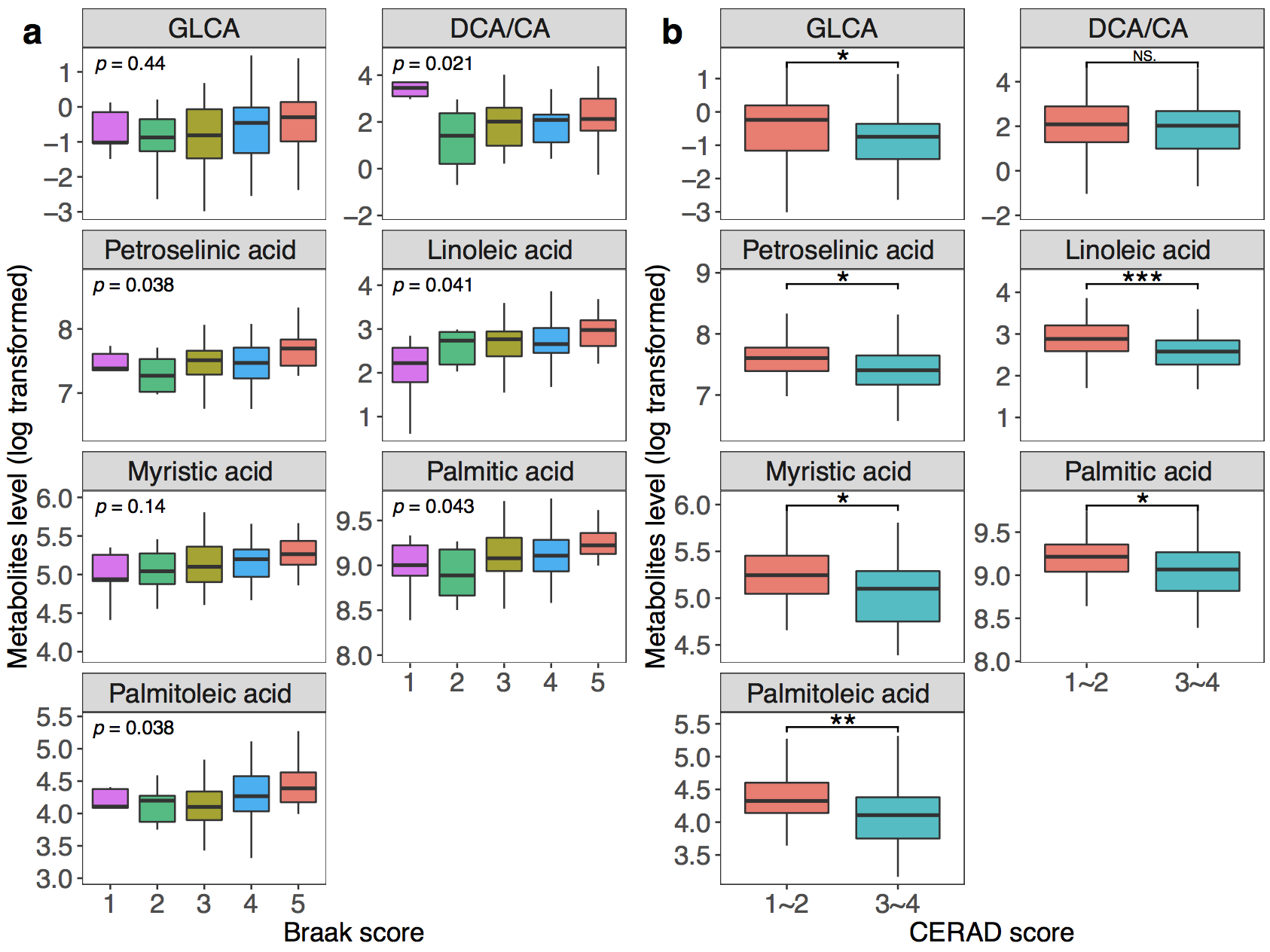

**Figure S8. Associations between metabolites level and Braak scores, CERAD scores.**

(a) Boxplots showing group differences and *P* values for identified metabolites across Braak groups for brain tissue abundances. (b) Boxplots showing group differences and significances for identified metabolites across CERAD groups for brain tissue abundances.

* *P*-value < 0.05, ** *P*-value < 0.01, *** *P*-value < 0.001, Wilcoxon rank sum test.

CERAD, Consortium to Establish a Registry for Alzheimer’s Disease; NS, not significant.

**Supplementary Text**

*Serum sample* *preparation*

We mixed an aliquot of 50 μL serum with 150 µL methanol, and vortexed the mixture for 2 min, let it stand for 10 min and centrifuged at 4 °C for 10 min. We then transferred 160 µL of the supernatant to a clean tube and vacuum dried the remaining aliquot re-dissolved it with a matched amount of acetonitrile (0.1% formic acid) and added water (0.1% formic acid) to a volume of 40 μL. The supernatant after the centrifugation was used for UPLC-TQMS and GC-TOFMS analysis. A mixture of 20 μL from the final supernatant of each sample was prepared for use as pooled quality control (QC) samples.

*Brain sample preparation*

We weighted 30 mg brain tissue, homogenized it with 75 µL of 50% precooled methanol using a Bullet Blender Tissue Homogenizer (Next Advance, Inc., Averill Park, NY) for 3 min. After centrifugation at 4 °C for 15 min, we transferred the supernatant to a clean tube. We performed the second step extraction by adding precooled methanol and chloroform mixture (3:1) to the residue followed by centrifugation. We combined the supernatant with the previous one and vortexed the mixture for 5 min and performed centrifugation for 15 min. The supernatant was used for UPLC-TQMS and GC-TOFMS analysis. A mixture of 20 μL from the final supernatant of each sample was prepared for use as pooled QC samples.

*Quality control Procedure*

Previously prepared QC samples were run between every ten sample injections. For each metabolite in QC samples, we calculated the relative standard deviations (RSDs), which was less than 15% for batches of samples.
